# Soft, 3D printed muscle ultrasound phantom with structurally tunable B-mode echo intensity

**DOI:** 10.1101/2024.11.29.625078

**Authors:** Samuel D Gillespie, Caralyn P Collins, Eric J Perreault, Cheng Sun, Oluwaseyi Balogun, Wendy M Murray

## Abstract

**OBJECTIVES:** Imaging phantoms for training and validation are vital to improving the performance and adoption of ultrasound imaging modalities in clinical and pre-clinical applications, and the goal of this study was to assess the viability of 3D printed muscle ultrasound phantoms to meet this need.

**METHODS:** We used a soft stereolithography resin to 3D print phantoms that mimicked the fascicle- and perimysium-scale structure of skeletal muscle and compared the long axis B-mode imaging quality and pattern of the phantom to that of healthy, adult Biceps brachii. We used a pulse-echo, time-of-flight method to measure the acoustic impedance of the resin for comparison to skeletal muscle and common soft tissue mimicking materials. We analyzed the echo intensity (EI) of muscle images to establish a physiological range and compared the EI of different phantom designs to assess the ability to control imaging brightness through structural modification.

**RESULTS:** A linear, striated hyper-/hypo-echoic B-mode imaging pattern mimicking long axis Biceps brachii muscle images was achieved with two 3D structure paradigms, rod and honeycomb. Acoustic impedance of Elastic 50A resin is higher than skeletal muscle in bulk, but appears suitable for use in a 3D structured phantom. EI measured in the Biceps images were found to vary both within and across images with an overall mean ± SD of 87 ±13 AU. EI measured in honeycomb phantoms (55 ±15 AU) was higher than in rod phantoms (42 ±13 AU), and a latticed honeycomb further increased EI (90 ±11 AU).

**CONCLUSIONS:** This study serves as proof-of-concept for soft, 3D printed phantoms that replicate the characteristic muscle ultrasound imaging pattern with the ability to tune clinically relevant EI values via structural design.

## INTRODUCTION

Due to the structure-function relationship of skeletal muscle^1-4^, measurement of macroscopic structural parameters (e.g., volume, cross-sectional area, fascicle length, pennation angle) provides fundamental information for the assessment of muscle function. In addition, skeletal muscle composition, characterized by the degree of adipose and/or fibrous tissue infiltration into muscle, provides an evaluation of muscle quality in healthy, diseased, and aging populations^5-8^. Magnetic resonance imaging (MRI) and computed tomography (CT) are the gold standard modalities for in-vivo imaging of skeletal muscle structure and composition^9^ but each suffers from limited accessibility^10, 11^ For CT, exposure to ionizing radiation is an additional consideration^12, 13^. B-mode ultrasonography is an accessible, safe, portable, non-invasive, and cost-effective alternative to evaluate skeletal muscle structure and composition^14^.

In-vivo ultrasound images of muscle structure are characterized by hypoechoic muscle fascicles surrounded by hyperechoic fibrous and adipose tissues^15^. The parallel arrangement of muscle fascicles results in a striped appearance in longitudinal images, and a spotted, “starry night” appearance in transverse images^16^. Clinically, these characteristic B-mode ultrasound imaging patterns allow for qualitative evaluations of muscle tissue (e.g., healthy, aging, spastic)^17^, diagnosis of musculoskeletal injury^15, 16^, and guidance for biopsies and injection-based therapy^18, 19^. The 4-point Heckmatt grading scale is a qualitative clinical assessment in which spastic muscle is graded visually based on its relative echo intensity (EI) compared to bone or fat; higher scores indicate a brighter image, suggesting reduced contractile material. When used in the clinical treatment of motor impairments for stroke, a higher Heckmatt score reflects a reduction in the potential therapeutic effect of Botox injection^17, 20^. EI can be quantified by measuring the mean pixel intensity within a specified region of interest; the interpretation of higher EI as an indication of increased non-contractile tissue infiltration within the muscle is supported by MRI, CT, and histological evidence^6, 14, 21-24^.

There is a critical, unmet need to improve validation methods for both clinical and pre-clinical research applications of musculoskeletal ultrasound. Improving validation of quantitative measures of muscle structural parameters using B-mode ultrasound commonly reported in research studies (e.g. fascicle length, pennation angle) has been identified as an important objective for the field^25^. Clinically, the use of ultrasound is highly user-dependent; a lack of standardized training limits the reliability of ultrasound assessments^26, 27^. Imaging phantoms provide a useful approach to evaluate, tune, and improve performance of any medical imaging modality. Current ultrasound imaging phantoms are comprised of molded tissue mimicking materials which replicate the bulk acoustic properties of human soft tissues^28^. The heterogeneous and anisotropic nature of skeletal muscle requires a phantom that can replicate not only bulk acoustic properties, but muscle’s characteristic imaging patterns that are dependent upon imaging plane, muscle structure and composition.

Development of muscle-like, B-mode imaging phantoms would serve as a first step toward addressing validation needs, as well as reliability and standardization of training in both research and clinical settings. We hypothesized that soft, 3D printed phantoms could mimic the structural geometry and ultrasound imaging properties of skeletal muscle, while also enabling rapid prototyping and customization of phantom design. We tested this hypothesis by designing and printing 2 phantom structures, one in which the printed structure mimicked the 3D geometry and dimensions of skeletal muscle fascicles (rod) and an inverted design intended to mimic the perimysium (honeycomb). We first qualitatively compared the B-mode imaging pattern of each design to in-vivo muscle images and measured the bulk acoustic properties (impedance and speed of sound) of the 3D printed resin for insight into the echogenicity of our phantoms and comparison to skeletal muscle and common tissue mimicking materials. We also quantified the echo intensity of B-mode images of our phantoms and directly compared to muscle images. Finally, we assessed our ability to adjust echo intensity of the phantom images via introduction of latticing to the honeycomb structure.

## MATERIALS

Stereolithography (SLA) printing was chosen for its capacity to 3D print high fidelity small features on the scale of muscle fascicles (100µ minimum feature size). FormLabs Elastic 50A resin (FormLabs, Somerville, MA) was selected as the softest commercially available SLA printer resin with a print area (145mm × 145mm × 185mm) that would allow for printing structures on the scale of whole human skeletal muscles. 3D models of muscle fascicle structure developed in SolidWorks 3D CAD software (Dassault Systemes, Vélizy-Villacoublay, France) were printed using the FormLabs Form3B SLA 3D printer. B-mode images of printed phantoms were recorded with a SIEMENS ACUSON S2000 ultrasound instrument using an 18L6 linear transducer. Long axis extended field-of-view (eFOV) ultrasound images of the Biceps brachii muscles of both arms of 4 healthy adult subjects acquired using the same ultrasound machine, obtained from a previous study^29^, were analyzed to compare the B-mode imaging pattern and brightness of phantom and in-vivo muscle images. Acoustic properties of the resin were quantified using a single GE 9LD wide band linear transducer with the Verasonics Vantage 256 Ultrasound Imaging System (Verasonics, Kirkland, WA).

## METHODS

### 3D Model Design

Human skeletal muscle fascicles are composed of 20-80 muscle fibers^30, 31^, each 20-100µm in diameter^32, 33^, grouped together and encapsulated by the perimysium, providing an estimated fascicle diameter range of 0.4-8mm. Using Solidworks 3D CAD software, we designed models to mimic the tissue structure of parallel fibered muscles on the fascicle and perimysium scale (Fig. 1). The rod design directly represents fascicles as solid, 50mm long, 2mm diameter printed fibers with unfilled space between them representing the perimysium. The honeycomb design inverts the rod design, with fascicles represented as unfilled, hexagonal pores of 2.17mm inscribed diameter in a 50mm long honeycomb-like printed structure representing the perimysium.

**Figure 1.**
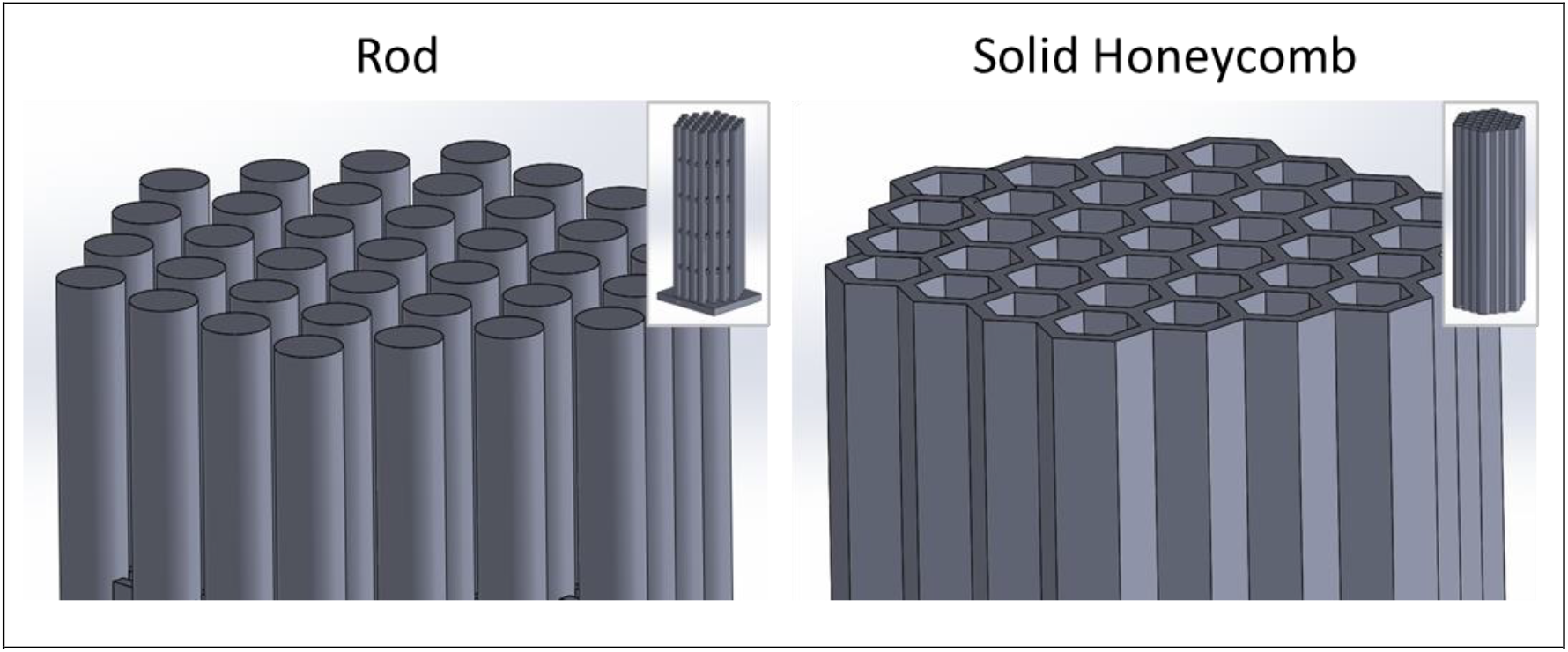
3D CAD models Magnified view of rod and honeycomb CAD models (Insets: full models).

### 3D Printing and Postprocessing

CAD models were imported as .STL files into FormLabs’ PreForm software for printing, and external supports were autogenerated by the software to enhance print stability. Using PreForm, the prepared models were then submitted to the printer. A Form3B SLA 3D printer was used to print the models using default settings for the Elastic 50A resin with a layer thickness of 100µm. In SLA 3D printing, the print platform, and subsequent cured layers, are lowered into a photocurable resin, and a rastering laser selectively cures a region as defined by the CAD model to form the next layer (Fig. 2). Upon print completion, external supports were removed and models were washed in isopropanol to remove uncured resin. The resulting structures were air dried and finished by curing under 405nm UV light at 60°C in Formlabs’ Form Cure mechanism for 20 minutes according to suggested curing conditions from Formlabs.

**Figure 2.**
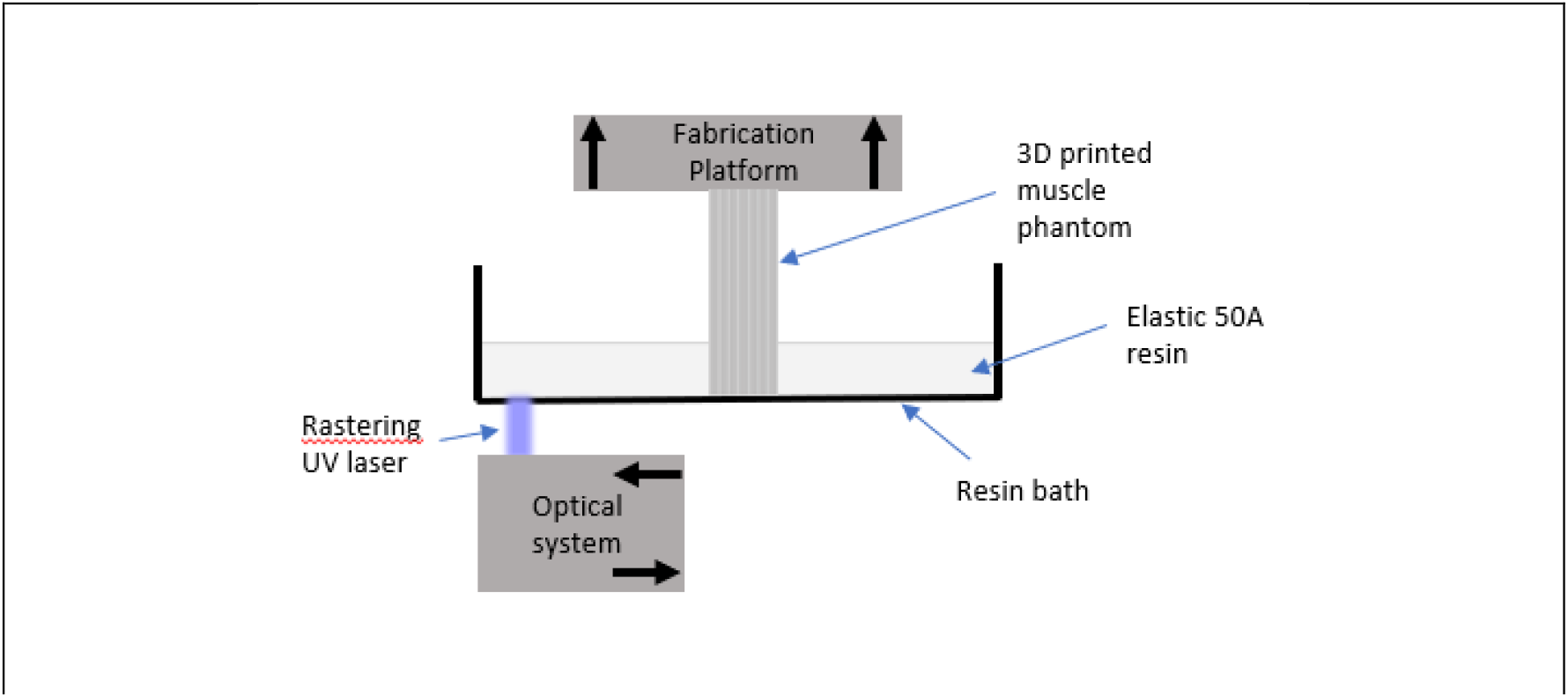
SLA 3D printing diagram. Diagram of SLA 3D printing process for muscle phantoms.

### Ultrasound Imaging

To evaluate the image quality of the 3D printed muscle phantoms, B-mode images were recorded with the Siemens Acuson 2000. Imaging parameters were set to the musculoskeletal exam default with 0dB gain and a frequency of 7MHz. Phantoms were submerged in water and images were recorded with the transducer in a vertical, long axis imaging plane (Fig. 3).

**Figure 3.**
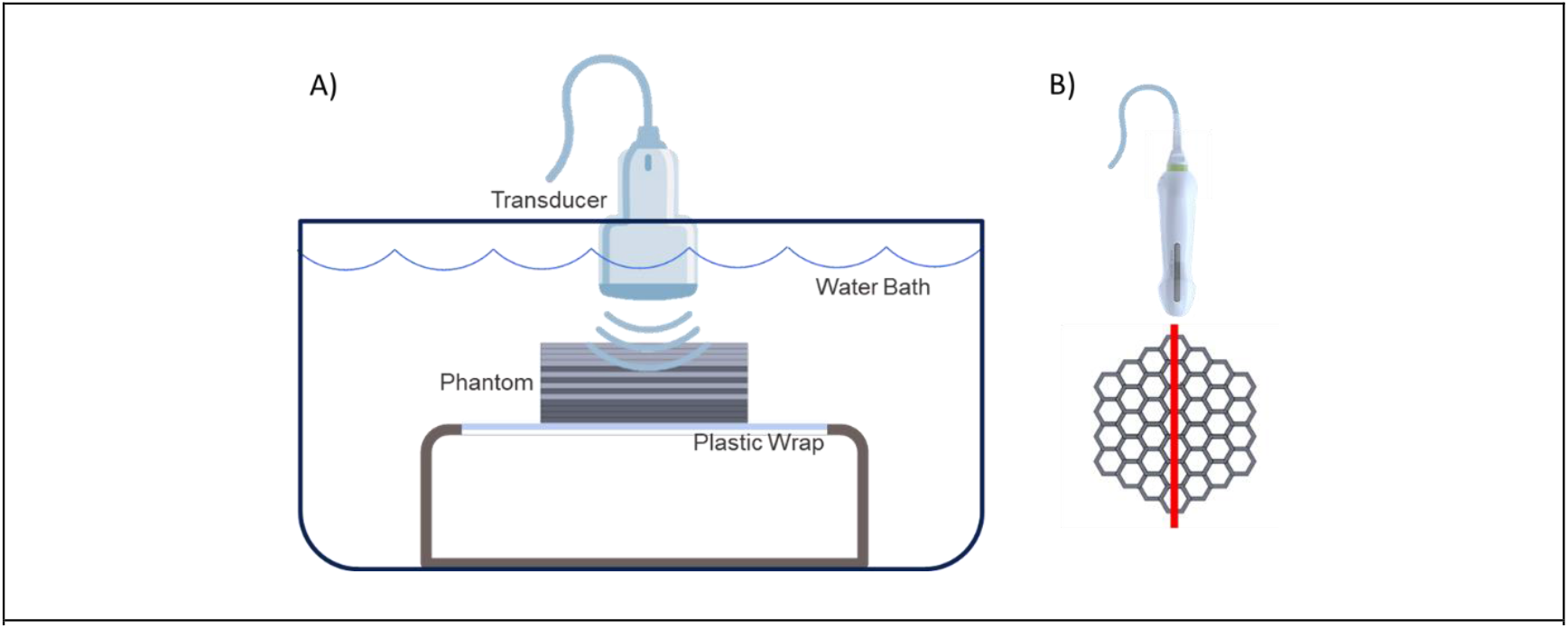
B-mode imaging set-up and transducer orientation. A) Diagram of phantom B-mode imaging set-up. B) Transverse illustration of the phantom and ultrasound transducer, with the red line indicating the imaging plane.

### Acoustic Testing

In order to test whether the acoustic properties of the selected resin were appropriate for skeletal muscle, we developed a modified pulse-echo, time-of-flight technique (Fig. 4) based upon previously reported methods used to quantify Speed of Sound (SoS) in TMMs^34-36^. Flat, rectangular samples (7cm × 3cm) with two different thicknesses (either 4mm or 5mm) were designed, printed, and post-processed. The resulting sample thickness (c.f., d, Fig. 4A) was confirmed before testing. In each measurement, a sample was secured to a reflective graphite wafer with the transducer fixed vertically to ensure signal propagation perpendicular to the plane of the sample and bottom surface. The transducer, sample, and reflective surface setup were immersed in a water bath for testing. A single transducer element was programmed to produce a sinusoid burst waveform with center frequency of 5.2083 MHz. The same transducer element remained active to receive the waveform reflections from the top surface of the sample and the bottom surface to which it was secured. The time point of the maximum absolute value of the transducer element voltage within the first reflected waveform (cf., *t*_1_, Fig. 4B) was subtracted from the corresponding time point of the second reflected waveform signal (c.f., *t*_2_, Fig. 4B) to determine the time-of-flight difference between the waves that reflected off the sample surface and those transmitted through the sample and reflected off the bottom surface. SoS was calculated as:

**Figure 4.**
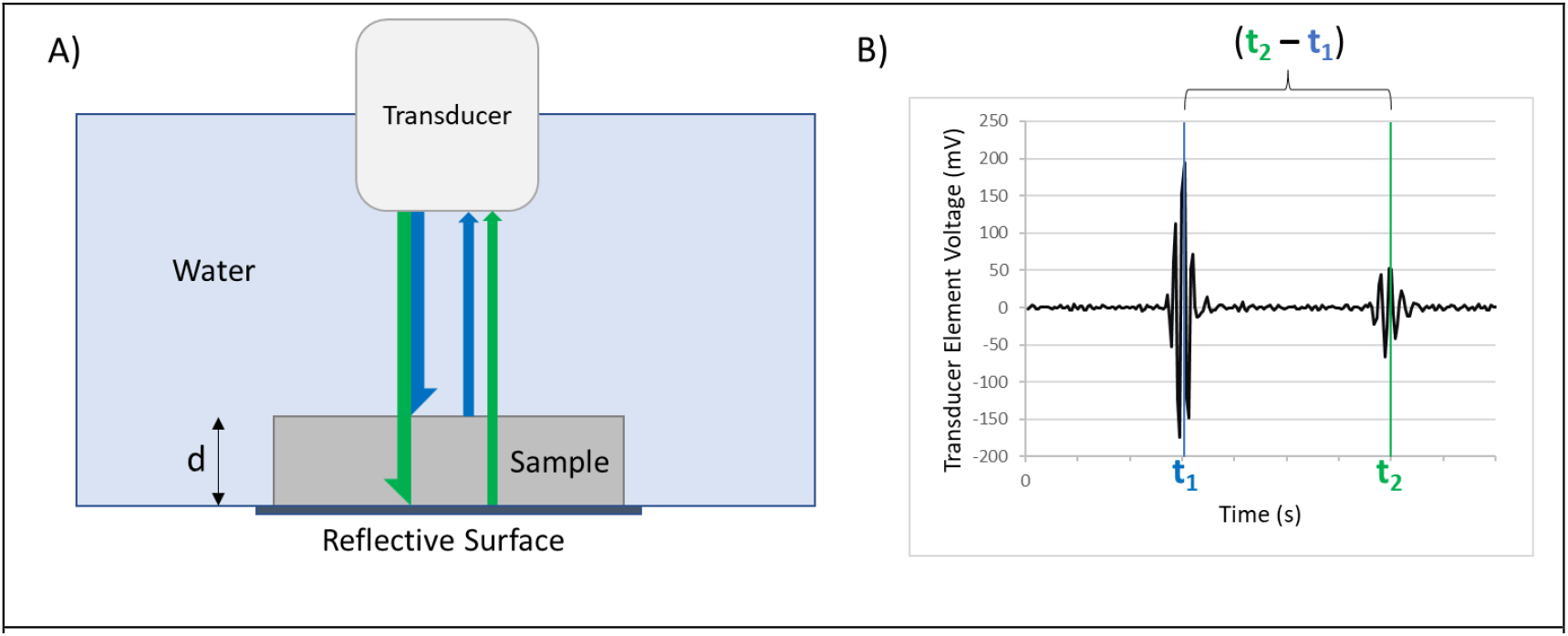
Modified pulse-echo, time-of-flight acoustic testing. **A)** Diagram illustrating the testing method used to assess bulk acoustic properties of the selected resin. Propagation path lengths of the waves reflected off the top and bottom surfaces are indicated by pairs of blue and green arrows, respectively. **B)** Representative graph of single element transducer voltage signal (mV) vs. time (seconds), showing the waveforms of the reflected signals. The signal that reflects off the top surface of the resin sample takes a shorter time (t_1_) to return compared to the time to return for the bottom reflected signal (t_2_).

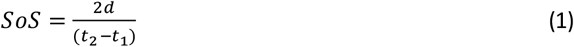

Specific acoustic impedance, *z*, of the resin was then calculated as the product of sample SoS and density, *ρ*:

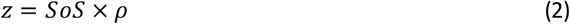

Sample density was determined separately by measuring the mass and volume of printed and cured cubes of resin. For method validation, measurement of the SoS in water with no sample present resulted in a value of 1485 ±3 ms^-1^ which agrees with the known value of 1482 ms^-1^ at 20°C^37, 38^.

### Echo Intensity Analysis

To assess the echo intensity (EI) levels of B-mode images of our 3D printed muscle phantoms relative to muscle images, EI analysis was performed of phantom and muscle images collected with the same ultrasound machine. All image analysis was completed with ImageJ (NIH, Bethesda, MD). To provide an in-vivo muscle EI baseline and relevant range, a total of 24 eFOV long axis Biceps brachii images were analyzed, comprised of three images acquired in both arms of four subjects^29^. Three rectangular regions of interest (ROIs) were defined within a single muscle in each image, characterizing image brightness in the distal, middle, and proximal regions of the image (Fig. 5). Four long axis phantom images were analyzed for each 3D printed phantom. Three rectangular ROIs were defined within each image, representing shallow, middle, and deep regions of the image. In all cases, B-mode ultrasound images saved in DICOM format were converted to 8-bit grayscale. EI within a selected ROI was defined as the average pixel intensity from 0 to 255 arbitrary units (AU).

**Figure 5.**
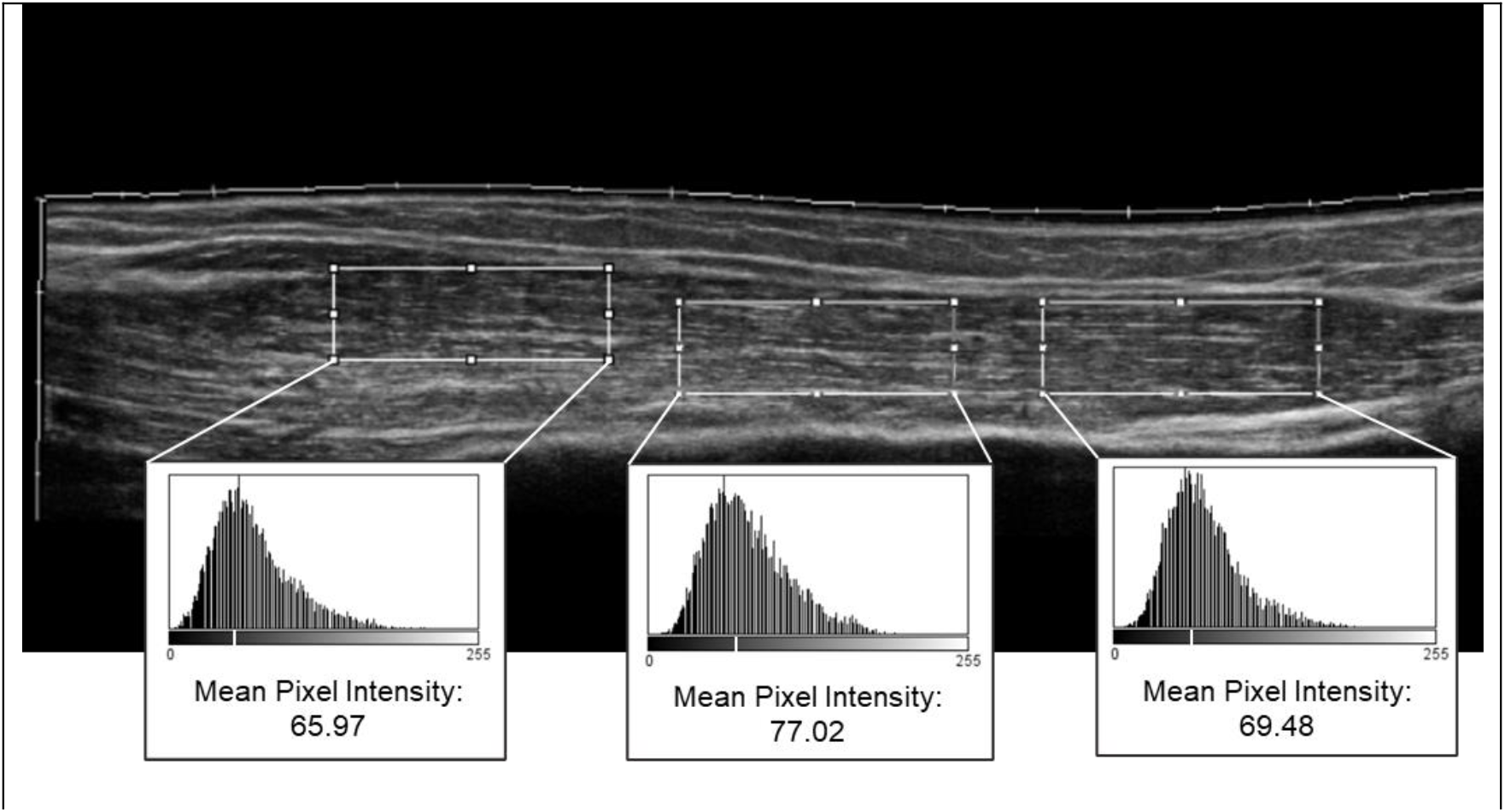
Biceps ROI and pixel intensity histograms. Long axis extended field-of-view Biceps brachii image with (left to right) Distal, Middle, and Proximal regions of interest outlined and corresponding grayscale pixel intensity histograms included. Echo intensity within each ROI is the mean pixel intensity value shown below the histogram.

### Echo Intensity Control

Given the clinical importance of alterations in echo intensity for assessment of muscle quality^17, 20, 39^, we evaluated the ability to adjust echo intensity of B-mode images of a phantom made from a specific resin via altering the 3D printed structure. To accomplish this, we introduced latticing to the honeycomb structure, with the intention of increasing ultrasound penetration and altering the reflection pattern of the resulting phantom. For this proof of concept analysis, we created a lattice design which introduces 1.25mm × 0.5mm gaps spaced 0.5mm apart in the long axis of the walls of each pore of the solid honeycomb design (Fig. 6).

**Figure 6.**
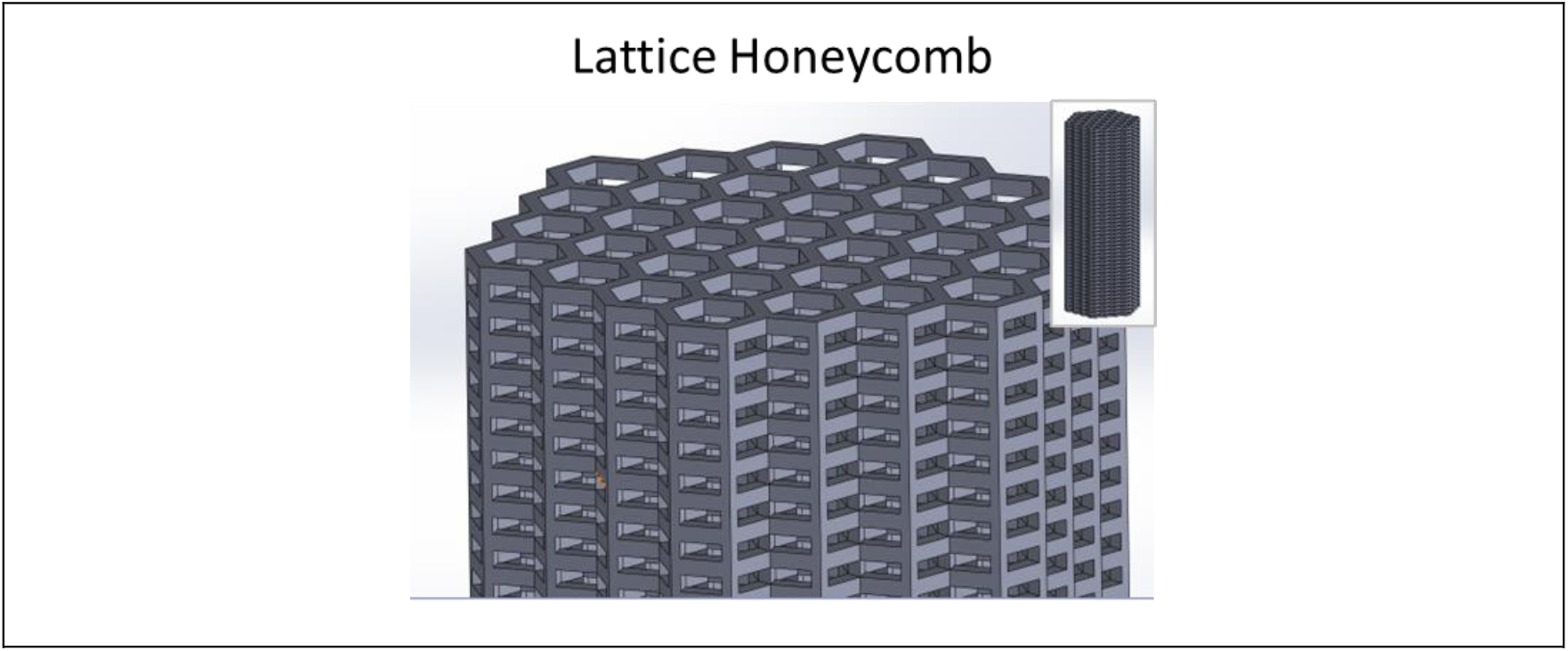
Lattice honeycomb 3D CAD model. Magnified view of Lattice Honeycomb CAD model (Inset: full model).

## RESULTS

### B-mode Pattern Replication

A linear, striated, alternating hyper- and hypo-echoic pattern mimicking long axis muscle B-mode images was achieved with both the 3D structure that mimicked the 3D geometry and dimensions of skeletal muscle fascicles (rod) and inverted design intended to mimic the perimysium (honeycomb). The linearly patterned geometry of both printed phantoms results in a highly regular contrast pattern that was deemed qualitatively successful (Fig. 7), despite its regularity, which lacks the natural variation found in muscle images.

**Figure 7.**
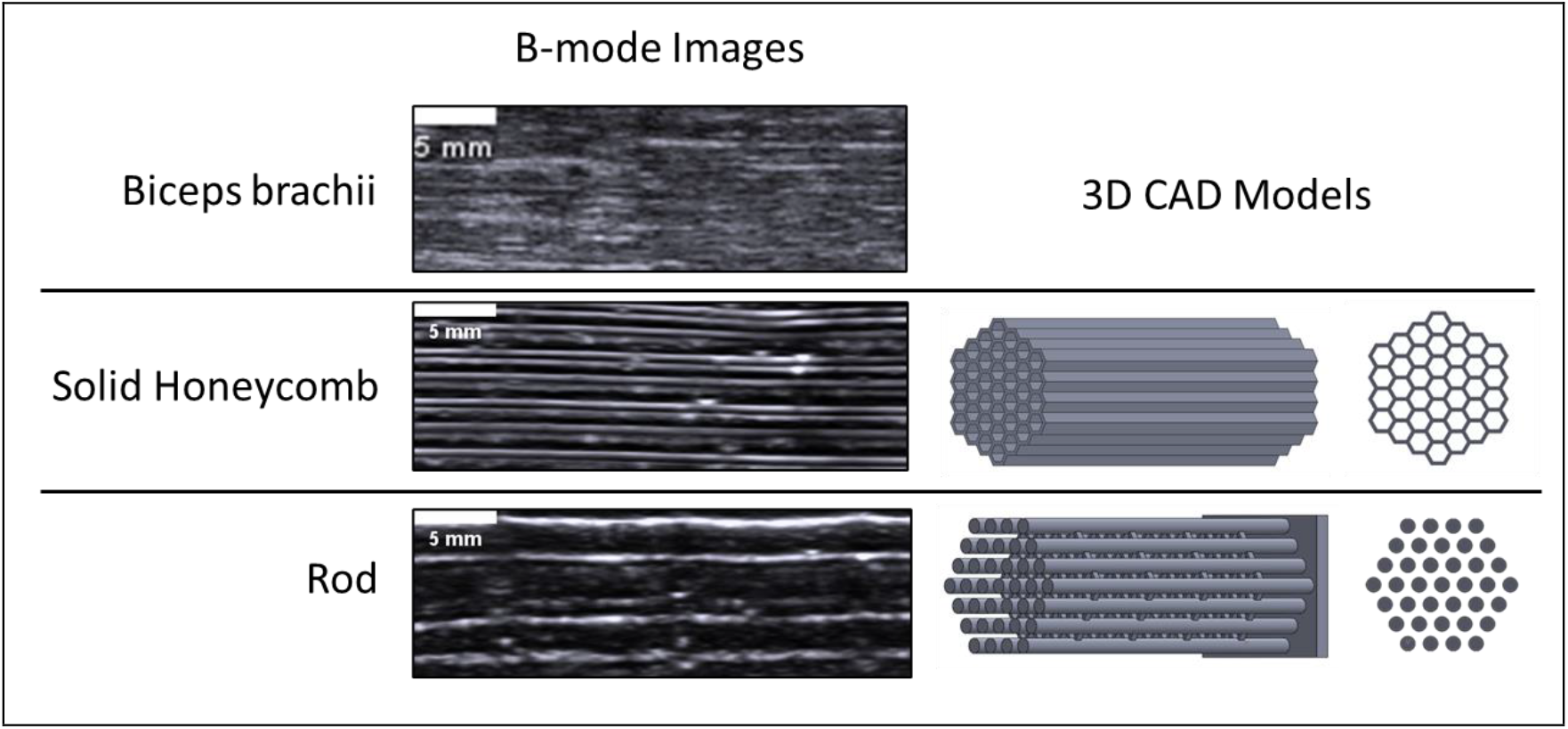
B-mode pattern replication. Representative long axis B-mode images of Biceps brachii muscle, honeycomb phantom, and rod phantom. Longitudinal and cross-sectional CAD models of honeycomb and rod phantoms.

### Acoustic Properties

The speed of sound within and acoustic impedance of Elastic 50A are higher than skeletal muscle and common bulk soft tissue mimicking materials. The measured ultrasonic speed of sound and calculated acoustic impedance of Elastic 50A are, respectively, 8% and 13% greater than the values reported for human skeletal muscle (Tbl. 1). For comparison, the SoS in Agarose, PAA, and PVA, respectively, are -5%, 0%, and -1% different and the corresponding impedance values are -5%, 4%, and 5% different than skeletal muscle.

**Table 1.** Acoustic properties of Elastic 50A, skeletal muscle, and common tissue mimicking materials. ^a^Standard deviations and measurement frequency not reported in skeletal muscle references

| Material | Speed of Sound<br>(m s<sup>-1</sup>) | SoS %<br>Difference | Impedance<br>(MRayl) | Impedance %<br>Difference | Frequency<br>(MHz) |
| --- | --- | --- | --- | --- | --- |
| Elastic 50A | 1710 ± 30 | 8% | 1.87 ± 0.03 | 13% | 5.2083 |
| <sup>a</sup>Skeletal Muscle<sup>40, 41</sup> | 1580 | - | 1.66 | - | - |
| Agarose 2%<sup>42</sup> | 1500 ± 30 | -5% | 1.57 ± 0.08 | -5% | 5 |
| PAA 10%<sup>42</sup> | 1580 ± 50 | 0% | 1.73 ± 0.08 | 4% | 5 |
| PVA<sup>42</sup> | 1570 ± 20 | -1% | 1.74 ± 0.08 | 5% | 5 |

**Table 1.** Acoustic properties of Elastic 50A, skeletal muscle, and common TMMs <sup>a</sup>Standard deviation and measurement frequency not reported in skeletal muscle reference
| Material | Speed of Sound<br>(m s<sup>-1</sup>) | SoS %<br>Difference | Impedance<br>(MRayl) | Impedance %<br>Difference | Frequency<br>(MHz) |
| --- | --- | --- | --- | --- | --- |
| Elastic 50A | 1710 ± 30 | 8% | 1.87 ± 0.03 | 13% | 5.2083 |
| <sup>a</sup>Skeletal Muscle<sup>40, 41</sup> | 1580 | - | 1.66 | - | - |
| Agarose 2%<sup>42</sup> | 1500 ± 30 | -5% | 1.57 ± 0.08 | -5% | 5 |
| PAA 10%<sup>42</sup> | 1580 ± 50 | 0% | 1.73 ± 0.08 | 4% | 5 |
| PVA<sup>42</sup> | 1570 ± 20 | -1% | 1.74 ± 0.08 | 5% | 5 |

### Muscle Echo Intensity Analysis

Within the 24 long axis eFOV Biceps brachii images we analyzed, there was echo intensity (EI) variation across all subsets of data, even within the same muscle in a single image. The overall mean EI value across all 72 ROIs in all images was 87 ±13 AU. The mean EI values from all 72 ROIs had an approximately normal distribution, spanning a range of 49 AU to 116 AU (Fig. 8B). The mean standard deviation of EI values within data subsets were; 4.3 AU within individual images, 6.7 AU within images of the same muscle, 7.5 AU within both arms of the same subject, and 12.8 AU across all data (Fig. 8C).

**Figure 8.**
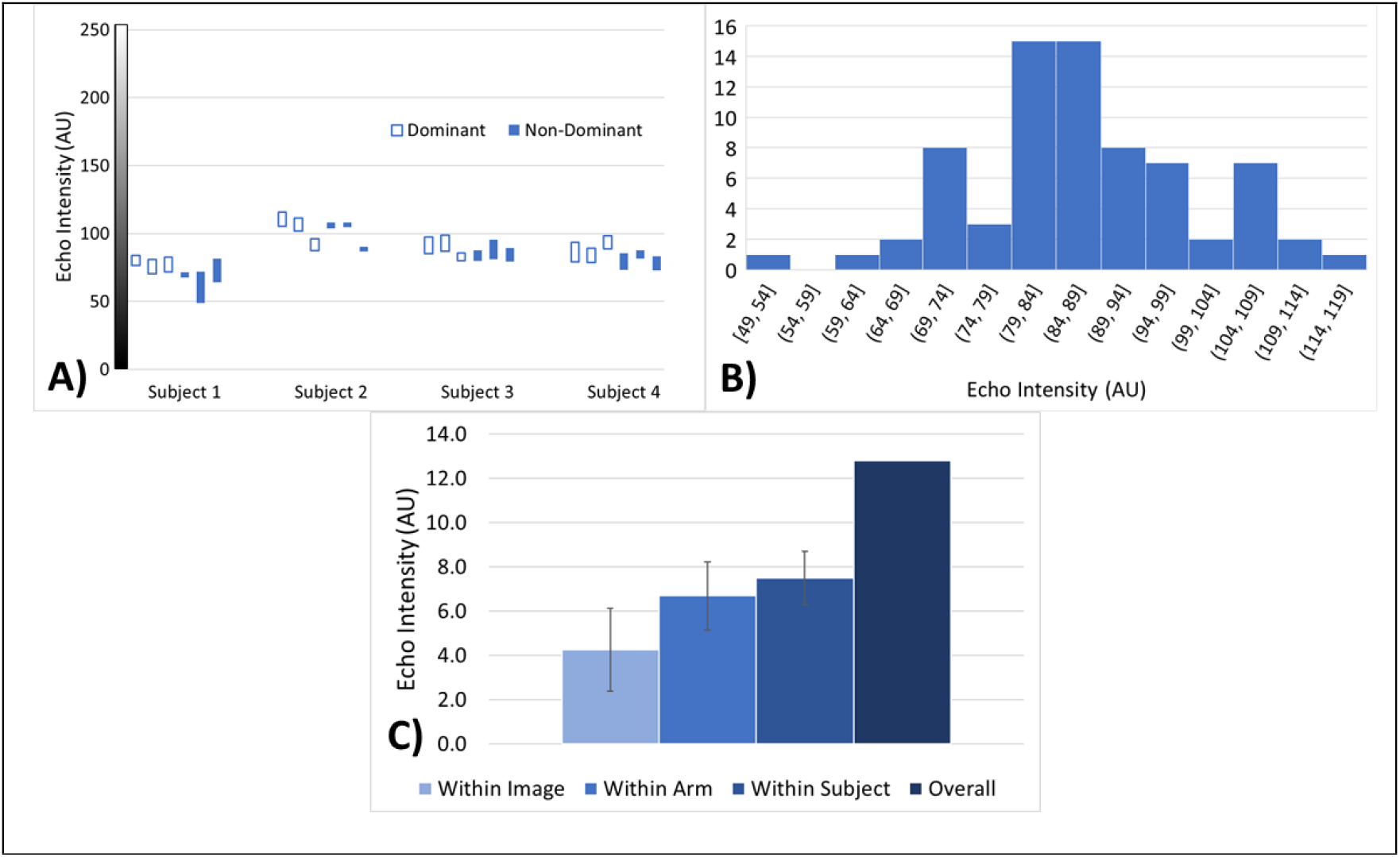
Muscle echo intensity analysis. **A)** Echo intensity (EI) range recorded in each of the 24 images plotted as bars. Data is grouped by subject (6 images per subject) with fill denoting arm dominance. **B)** Histogram of EI values recorded in all 72 ROI across the 24 images. **C)** Mean standard deviation of EI within different subsets of data; within the same image, within all images of the same arm, within all images of the same subject (both arms), and overall across all images.

### Phantom Echo Intensity Control

In phantoms printed using the same resin, B-mode echo intensity levels varied depending on the 3D structure. Echo intensity measured in the solid honeycomb was higher than the rod, and the lattice honeycomb design further increased the EI and most closely replicated the overall mean EI measured in long axis Biceps brachii images. Echo intensity values (mean ± SD) measured in phantom B-mode images of the rod, solid honeycomb, and lattice honeycomb designs were 42 ±13 AU, 55 ±15 AU and 90 ±11 AU, respectively (Fig. 9).

**Figure 9.**
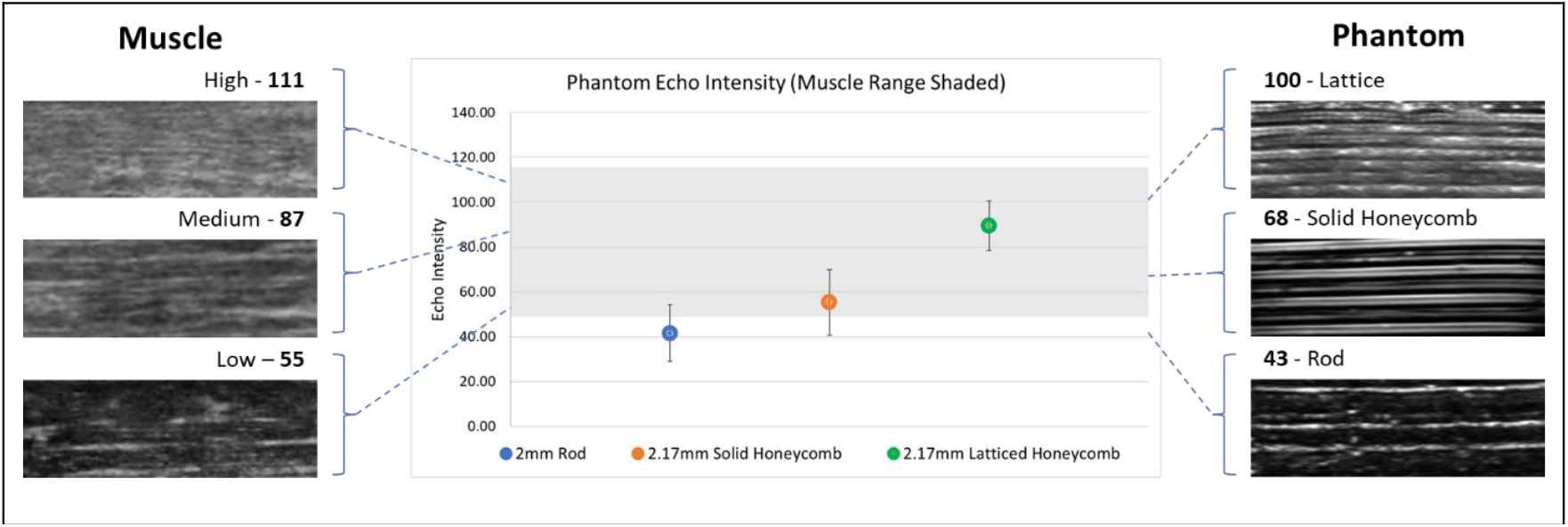
Phantom echo intensity control. (Center) Plot of echo intensity (EI) values (Mean ± Standard Deviation) of rod, solid honeycomb, and lattice honeycomb designs. Grey area indicates range of EI values recorded in long axis Biceps brachii images. (Left) Representative high, medium, and low EI regions within muscle B-mode images. (Right) Representative lattice honeycomb, solid honeycomb, and rod phantom B-mode images.

## DISCUSSION

In-vivo muscle ultrasound is increasingly used to study muscle structure and composition in clinical and pre-clinical settings, but there remains a critical need for validation and training methods to improve adoption and reliability of ultrasound assessments. Our goal was to investigate the feasibility of using a soft, elastomeric resin to create 3D printed phantoms that mimic the acoustic and ultrasound imaging qualities of human skeletal muscle. We designed 3D structures mimicking muscle fascicles and perimysium and assessed their B-mode imaging pattern compared to in-vivo muscle images. We then tested the speed of sound and acoustic impedance of the soft, elastomeric SLA resin for insight into the echogenicity of our phantoms. Finally, we analyzed the echo intensity of our phantoms and introduced latticing to our design to demonstrate our ability to control phantom echo intensity through structural changes.

The 3D printed phantoms based upon skeletal muscle fascicle and perimysium dimensions successfully demonstrated an ability to mimic the striped hyper- and hypo-echoic pattern of in-vivo images. The hyperechoic lines in the phantoms are created by reflection from the material surfaces of the fibers and pores within the rod and honeycomb structures, respectively, and the hypoechoic spacing corresponded to the internal gaps within the phantom that were filled with water when submerged during imaging. A comparison of the hyperechoic line spacing in the Biceps brachii images, measured as low as 0.2mm, and the phantom images, measured no lower than 0.5mm as expected given design dimensions, indicates a need to reduce the printed structure dimensions to match in-vivo muscle images more closely. Initial efforts to reduce the fiber and pore diameters to match the dimensions of muscle imaging patterns identified fiber print stability (rod) and pore resin drainage (honeycomb) as issues that prevented designs with diameters below ∼2mm. Even with 2mm fibers, rod phantoms required the use of supports between fibers to maintain stability during printing. The introduction of latticing to the honeycomb holds promise for not only echo intensity tuning, but also to allow for increased resin drainage and further pore diameter reduction. Assessment of the variability of hyperechoic spacing in different muscles and subjects would help to set a dimensional goal for pore/fiber diameter and define the design parameter changes necessary for muscle-like phantoms to mimic specific muscles.

The measured speed of sound and acoustic impedance of Elastic 50A resin was higher than literature values for skeletal muscle and other commonly adopted soft tissue mimicking materials. These soft tissue mimicking materials are prepared via molding and therefore lack the capacity to control internal 3D structure, a major advantage of 3D printing^28, 43^. As opposed to molded materials which mimic the bulk acoustic properties of skeletal muscle, our phantoms introduce a 3D structure. As such, the properties of the resin alone do not accurately represent the bulk properties of a printed phantom. The higher acoustic impedance of the printed resin places surface reflection as the predominant mode of echogenic contrast within the phantom, as opposed to the volumetric scattering of muscle tissue and bulk tissue mimicking materials. In combination with high fidelity printing of fascicle-scale 3D structures, this mode of reflection can be harnessed to mimic the anisotropic nature and imaging pattern of skeletal muscle. Further work needs to be done to characterize the bulk acoustic properties of 3D patterned phantoms immersed in water or backfilled with a soft material.

Due to the periodic geometry of the printed phantoms, image quality and brightness were dependent upon the imaging plane. All images shown in the study and used for echo intensity analysis were recorded in the vertical, long axis plane of the phantoms which provided the highest quality images and greatest hyper- and hypo-echoic contrast. The high user dependence of clinical muscle imaging, which involves determination of an appropriate imaging plane, serves as an analogy and potential application of this angle dependent imaging in training new clinicians. The periodic geometry of the phantoms also creates the unnaturally regular B-mode imaging pattern because the hyperechoic lines are generated by perfectly straight, regularly spaced material surfaces. Incorporating realistic muscle cross-sectional patterns and fascicle paths via translation of in-vivo muscle images^44-46^ or muscle finite element modelling strategies^47^ to the CAD models could eliminate the highly periodic geometry and create imaging phantoms that more closely resemble the natural variation found in muscle.

Echo intensity is a useful and accessible tool for quantifying muscle quality via ultrasound, but there are many confounding factors that limit comparisons across different instruments, imaging parameters, and subjects. This variation found within and across in-vivo muscle B-mode images, and the fact that echo intensity differences as opposed to exact values are important for clinical evaluation, indicates that there is no correct, “gold-standard” echo intensity value that a muscle phantom should aim to replicate. It is more important that a phantom design has the tunability to replicate the range of echo intensity values that are likely to be found in muscle images from diverse subject groups (e.g., healthy, strained, aging, spastic). The introduction of latticing as a structural control of echo intensity and the ability to rapidly prototype and adjust design parameters is a major benefit of 3D printed phantoms. At the measurement frequency of 5.2083MHz, the wavelength of ultrasound waves is ∼0.3mm within water and resin mediums, which is only slightly shorter than the width of the smallest lattice features, 0.5mm. As the feature size of lattice structures approaches and drops below the wavelength of the ultrasound waves, reflection will more closely resemble the scattering/diffuse reflection and brightness observed in skeletal muscle as opposed to the specular reflection and stark, light/dark contrast observed in the solid walled honeycomb and rod designs. Investigation of the relationship between lattice design parameters and resulting echo intensity is needed to move toward clinically relevant muscle ultrasound phantoms from these proof-of-concept designs.

## CONCLUSION

The high-user dependency limiting the adoption need for muscle ultrasound phantoms to improve the validity and training methods of clinical and pre-clinical ultrasound applications. This would enable greater adoption of the imaging modality and enhance the quality of ultrasound assessments. SLA 3D printing with FormLabs’ Elastic 50A resin was investigated as a method to mimic the anisotropic structure and imaging pattern of skeletal muscle. The acoustic properties of the resin combined with the 3D structure of the printed phantoms enabled replication of the hyper- and hypo-echoic pattern, while introduction of latticing created a structural control of the echo intensity of phantom images. This study serves as proof-of-concept that soft, 3D printed phantoms can replicate the characteristic muscle ultrasound imaging pattern with the ability to tune physiologically relevant echo intensity values via structural design.

## ACKNOWLEDGMENTS

This work was supported by the Department of Veterans Affairs RR&D through the SPiRE I21 funding mechanism (Grant # I21RX003610).

## CONFLICT OF INTEREST STATEMENT

No conflicts of interest to report.

## DATA AVAILABILITY STATEMENT

Investigate/decide how to share research data

## Notes

### Competing Interest Statement

The authors have declared no competing interest.

